# No clear evidence for correlations between handgrip strength and sexually dimorphic acoustic properties of voices

**DOI:** 10.1101/227165

**Authors:** Chengyang Han, Hongyi Wang, Vanessa Fasolt, Amanda C Hahn, Iris J Holzleitner, Junpeng Lao, Lisa M DeBruine, David R Feinberg, Benedict C Jones

**Author notes:** **Corresponding author:** Chengyang Han, Institute of Neuroscience & Psychology, University of Glasgow, Scotland, UK. Data and analysis files are publicly available at https://osf.io/na6be/.

## Abstract

**Objectives:** Recent research on the signal value of masculine physical characteristics in men has focused on the possibility that such characteristics are valid cues of physical strength. However, evidence that sexually dimorphic vocal characteristics are correlated with physical strength is equivocal. Consequently, we undertook a further test for possible relationships between physical strength and masculine vocal characteristics.

**Methods:** We tested the putative relationships between White UK (N=115) and Chinese (N=106) participants’ handgrip strength (a widely used proxy for general upper-body strength) and five sexually dimorphic acoustic properties of voices: fundamental frequency (F0), fundamental frequency’s standard deviation (F0-SD), formant dispersion (Df), formant position (Pf), and estimated vocal-tract length (VTL).

**Results:** Analyses revealed no clear evidence that stronger individuals had more masculine voices.

**Conclusions:** Our results do not support the hypothesis that masculine vocal characteristics are a valid cue of physical strength.

## Introduction

Early research on the signal value of masculine physical characteristics in men focused on the possibility that masculine physical characteristics were valid cues of men’s immunocompetence (Penton-Voak et al., 1999; Thornhill & Gangestad, 1999). More recent research has focused on the possibility that masculine physical characteristics may, instead, be valid cues of men’s physical strength (reviewed in Puts, 2010 and Scott et al., 2013). Evidence supporting this latter hypothesis has come from studies reporting positive correlations between men’s facial masculinity and upper-body strength, as measured via handgrip strength (Fink et al., 2007; Windhanger et al., 2011). Further evidence for this hypothesis comes from research showing that increasing upper-body strength increases the masculinity of men’s upper-body shape (see Fan et al., 2005).

Although there is good evidence that more masculine male voices are perceived to be more dominant (e.g., Feinberg et al., 2005; Puts, 2006), only two studies have explored possible relationships between upper-body strength and sexually dimorphic vocal characteristics. Puts et al. (2012) investigated possible relationships between US and Hadza men’s arm strength (a composite measure derived from handgrip strength and flexed bicep circumference) and four sexually dimorphic vocal characteristics; fundamental frequency (F0), fundamental frequency’s standard deviation (F0-SD), formant dispersion (Df), and formant position (Pf). Values for F0, F0-SD, Df, and Pf are typically larger in women than men (see, e.g., Puts et al., 2012). The fundamental frequency (F0) is produced by the vibration of the vocal folds and perceived as voice pitch, whereas formants are the resonant frequencies of the vocal tract (Pisanski et al., 2014a, 2016). Stronger US men tended to have more masculine F0-SD and more masculine Pf. Stronger Hadza men tended to have more masculine F0 (see also Hodges-Simeon et al., 2014 for similar findings in circum-pubertal males). Df did not predict handgrip strength in either group. Although these correlations suggest men with more masculine voices may be physically stronger, the significant relationships would not have survived correction for multiple comparisons, suggesting they may not be robust (Puts et al., 2012). Indeed, Sell et al. (2010) found no significant relationships between physical strength and either F0 or Df in US, Tsimane, or Andean men, or in a sample of US women. Moreover, in a sample of Hadza men that overlapped with that analyzed in Puts et al. (2012), Smith et al. (in press) reported no significant correlation between upper-body strength and F0.

The current study reports a further test of the hypothesized relationship between upper-body strength and sexually dimorphic acoustic properties of voices in a sample including both White UK and Chinese men and women. While most research on this topic has investigated only men, here we investigated both men and women. Following Fink et al. (2007), Windhanger et al. (2011), and Puts et al. (2012), upper-body strength was assessed via handgrip strength. In addition to the F0, F0-SD, Pf, and Df measures considered by Puts et al. (2012), we measured a fifth sexually dimorphic vocal characteristic (estimated vocal-tract length, VTL, Reby & McComb, 2003). We included VTL because of meta-analytic evidence that it predicts body size in men and women and may, therefore, be related to strength (Pisanski et al., 2014a). Values for VTL are generally larger in men than women (see, e.g., Pisanski et al., 2014a).

## Methods

In total, 221 participants took part in the study. These included 58 White UK men and 57 White UK women, all of whom were born and resided in the UK. They also included 53 Chinese men and 53 Chinese women, all of whom were born in China, but currently resided in the UK (mean number of years in the UK = 1.04 years, SD = 0.93 years). Mean age (and SD) for each group of participants is given in Table 1. None of these participants had taken part in our previous research on vocal cues of men’s threat potential (Han et al., 2016).

We measured each participant’s handgrip strength from their dominant hand two times using a T. K. K. 5001 Grip A dynamometer. Following Fink et al. (2007), the highest recording from each participant (i.e., their maximal handgrip strength) was used in analyses (see Table 1 for means and SDs). A mono digital voice recording of each participant was also taken, using an Audio-Technica AT-4041 cardioid condenser microphone at a sampling rate of 44.1 kHz at 16-bit amplitude quantization. Each participant was instructed to say “Hi, I’m a student at the University of Glasgow” in their normal speaking voice. White UK participants were recorded speaking English and Chinese participants were recorded speaking Mandarin.

All acoustic measurements were made using Praat (Boersma & Weenink, 2013). F0 was measured using Praat’s autocorrelation algorithm with a search range of 100-600 Hz for women and 65-300 Hz for men (Pisanski et al., 2014a). We measured F1 to F4 using Praat’s Burg Linear Predictive Coding (LPC) algorithm, with the maximum formant set to 5000 Hz for men and 5500 Hz for women. Formants were first overlaid on a spectrogram and formant number was manually adjusted until the best visual fit of predicted onto observed formants was obtained. These techniques and settings are recommended by the Praat manual (Boersma & Weenink, 2013; see also Pisanski et al., 2014a). F0, F0-SD, Pf, and Df were calculated using methods described in Puts et al. (2011) and estimated VTL was also calculated from formant frequencies using a formula described in Reby and McComb (2003). Each variable’s means (and SDs) are shown in Table 1 for each group of participants. Independent samples t-tests showed that each of these acoustic measures was sexually dimorphic in both White UK (all absolute t>11.0, all p<.001) and Chinese (all absolute t> 10.8, all p<.001) speakers.

**Table 1.**
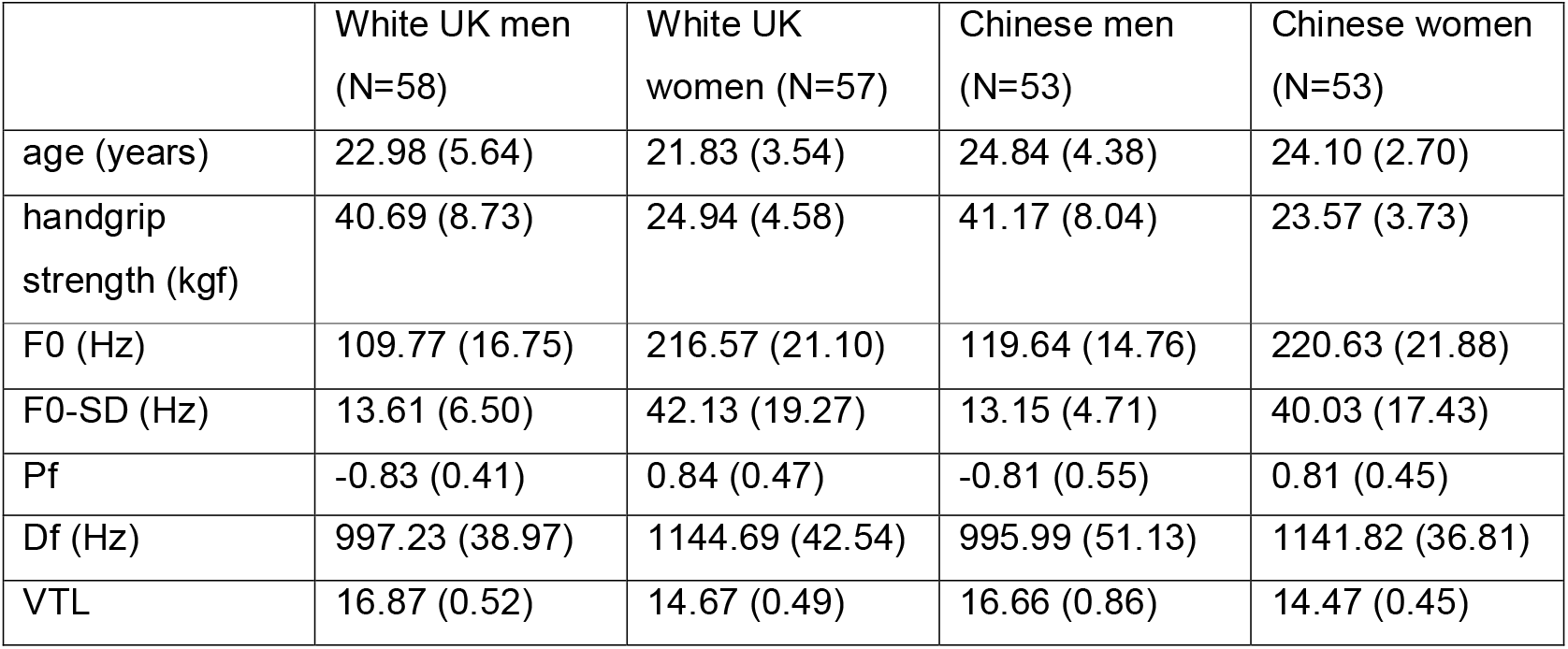
Each variable’s means (and SDs) for each group of participants.

|  | White UK men<br>(N=58) | White UK<br>women (N=57) | Chinese men<br>(N=53) | Chinese women<br>(N=53) |
| --- | --- | --- | --- | --- |
| age (years) | 22.98 (5.64) | 21.83 (3.54) | 24.84 (4.38) | 24.10 (2.70) |
| handgrip<br>strength (kgf) | 40.69 (8.73) | 24.94 (4.58) | 41.17 (8.04) | 23.57 (3.73) |
| F0 (Hz) | 109.77 (16.75) | 216.57 (21.10) | 119.64 (14.76) | 220.63 (21.88) |
| F0-SD (Hz) | 13.61 (6.50) | 42.13 (19.27) | 13.15 (4.71) | 40.03 (17.43) |
| Pf | -0.83 (0.41) | 0.84 (0.47) | -0.81 (0.55) | 0.81 (0.45) |
| Df (Hz) | 997.23 (38.97) | 1144.69 (42.54) | 995.99 (51.13) | 1141.82 (36.81) |
| VTL | 16.87 (0.52) | 14.67 (0.49) | 16.66 (0.86) | 14.47 (0.45) |

## Results

Analyses were conducted using R version 3.4.2 (R Core Team, 2016). Each of the five acoustic measures (F0, F0-SD, Pf, Df, VTL) was analyzed in a separate model. Handgrip strength was the dependent variable in each model. Predictors were acoustic measure (scaled and centered on the grand mean), speaker ethnicity (effect coded: Chinese = 0.5, White UK = −0.5), speaker sex (effect coded: male = 0.5, female = −0.5), and all possible interactions among these predictors. Full model specifications are given in our Supplemental Materials. Data files and analysis scripts are publicly available at https://osf.io/na6be/.

Each of the five models showed a significant effect of speaker sex (all estimates>12.90, all ts>4.90, all ps<.001), confirming that men had significantly greater handgrip strength than did women (see Table 1). No other main effects (all absolute ts<1.90, all ps>.07) or interactions (all absolute ts<1.90, all ps>.06) were significant in any of the models. None of the models showed effects of acoustic properties that were significant or approached significance (all absolute estimates<1.40, all absolute ts<1.50, all ps>.15). Full results for each model are given in our Supplemental Materials.

In addition to these regression analyses, we performed a Bayesian analysis with a multivariate latent model. The estimates of the correlations between handgrip strength and acoustic measures from this analysis overlapped with zero (bracketed numbers are the 95% highest posterior density). The correlation between handgrip strength and F0 was 0.145 [-0.407, 0.099]. The correlation between handgrip strength and F0-SD was 0.115 [-0.133, 0.365]. The correlation between handgrip strength and Df was −0.166 [-0.369, 0.055]. The correlation between handgrip strength and Pf was −0.101 [-0.303, 0.087]. The correlation between handgrip strength and VTL was 0.116 [-0.052, 0.294]. These results provide converging evidence that the sexually dimorphic acoustic characteristics of voices we measured were not reliably related to handgrip strength. Details of these analyses and full results are reported in our Supplemental Materials and at https://osf.io/na6be/.

## Discussion

We investigated hypothesized relationships between five sexually dimorphic acoustic properties (F0, F0-SD, Df, Pf, VTL) of voices and handgrip strength (a widely used proxy for upper-body strength). Our analyses revealed no clear evidence that stronger individuals had more masculine voices. These results are consistent with other work finding similar null results (Sell et al., 2011; Smith et al., in press), suggesting that previously reported positive correlations between arm strength and sexually dimorphic vocal characteristics (Puts et al., 2012) may not be robust. Our null results are also consistent with recent work suggesting that voices may not necessarily be a valid cue of men’s threat potential (see also Han et al., 2017).

Although we do not replicate the significant correlations reported previously between acoustic properties of voices and upper-body strength in circum-pubertal males (Hodges-Simeon et al., 2014) and men (Puts et al., 2012), this may be partly due to methodological differences. For example, those studies assessed strength using a measure that combined handgrip strength and bicep circumference. It is unlikely that we were underpowered to detect putative relationships between upper-body strength and vocal characteristics or that the absence of significant relationships between handgrip strength and vocal characteristics in our study is a consequence of not controlling for measures of body size. Our sample size of 111 men or 110 women gave us 80% power to detect simple correlations between vocal masculinity and strength with |r| > .23 at alpha = 0.05, which is slightly smaller than the significant zero-order correlations reported between upper-body strength and vocal characteristics in previous research reporting positive associations between vocal masculinity and strength (e.g. Puts et al. (2012): r = -.26 for 176 men). If we have failed to detect a reliable relationship between strength and vocal characteristics in the current study, we suggest it is likely to be due to how we measured strength or low variation in strength in our sample. Although the speech samples we used are similar to those employed in previous research on this topic, stronger relationships between strength and vocal characteristics might occur using less neutral vocalizations (e.g., the type of vocalizations made during direct competition), if individuals exaggerate existing (but otherwise subtle) vocal cues of strength when competing. Nonetheless, we note here that several studies have now found no significant correlations between strength and vocal masculinity in men (Sell et al., 2011; Smith et al., in press; Kordsmeyer et al., 2018) and the significant correlations reported by Puts et al. (2012) were both inconsistent across samples and would not have been significant if corrected for multiple comparisons. Collectively, these points may indicate the correlations reported in Puts et al. (2012) might be false positives).

If sexually dimorphic characteristics of voices are not reliably correlated with physical strength, what physical characteristics do they signal? One possibility is that they signal other aspects of threat potential. Consistent with this proposal, a recent meta-analysis found that some sexually dimorphic vocal characteristics (e.g., VTL) do reliably (although weakly) predict within-sex variation in adult body size (Pisanski et al., 2014a). Alternatively, associations between masculine vocal characteristics and threat-related perceptions (e.g., dominance) could be byproducts of sensory biases, such as a generalization of the tendency for larger objects to make lower-pitched sounds (Rendall et al., 2007; Pisanski et al., 2014b). Work exploring this latter possibility may prove fruitful.

In conclusion, our analyses showed no clear evidence for a positive association between masculine vocal characteristics and handgrip strength. These null results do not support the hypothesis that masculine vocal characteristics are a valid cue of physical strength.

## Author Contributions

CH, BJ, and DF designed the study. CH, HW, and VF collected the data. CH, BJ, LD, DF and JL analyzed the data. CH and BJ drafted the manuscript. AH and IH edited the manuscript for intellectual content and provided critical comments on the manuscript.

